# Force-dependent recruitment of Piezo1 drives adhesion maturation and calcium entry in normal but not tumor cells

**DOI:** 10.1101/2020.03.09.972307

**Authors:** Mingxi Yao, Ajay Tijore, Delfine Cheng, Jinyuan Vero Li, Anushya Hariharan, Boris Martinac, Guy Tran Van Nhieu, Charles D Cox, Michael Sheetz

**Affiliations:** Department of Biomedical Engineering, Southern University of Science and Technology; Guangdong Provincial Key Laboratory of Advanced Biomaterials; Mechanobiology Institute, National University of Singapore Mechanobiology Institute, National University of Singapore; Centre for Biosystems Science and Engineering, Indian Institute of Science, Bangalore, 560012, India; Victor Chang Cardiac Research Institute, Sydney, Australia; Ecole Normale Supérieure PARIS-SACLAY; Dept of Biological Sciences, National University of Singapore; Molecular MechanoMedicine Program, Dept. of Biochemistry and Molecular Biology, University of Texas Medical Branch

## Abstract

Mechanosensing is an integral part of many physiological processes including stem cell differentiation, fibrosis, and cancer progression. Two major mechanosensing systems – focal adhesions and mechanosensitive ion channels, can convert mechanical features of the microenvironment into biochemical signals. We report here surprisingly that the mechanosensitive Ca^2+^-channel Piezo1, previously perceived to be diffusive on plasma membranes, binds to matrix adhesions in a force-dependent manner, promoting adhesion maturation and cell spreading in normal but not in tumor cells. In the absence of Piezo1, matrix adhesions are smaller in normal cells mimicking transformed cells where adhesions do not change with or without Piezo1. A novel adhesion-targeted calcium sensor shows robust Piezo1-dependent, calcium influx at adhesions in normal cells; but not in transformed cells. A linker domain in Piezo1 is needed for binding to adhesions and overexpression of the domain blocks Piezo1 binding to adhesions decreasing adhesion size and cell spread area. Thus, we suggest that Piezo1 is a novel component of focal adhesions in non-transformed cells that catalyzes adhesion maturation and growth through force-dependent calcium signaling, but this function is absent in most cancer cells.

## Introduction

Cellular mechanosensing, the cell’s ability to detect and respond to its physical microenvironment, is involved in important processes in development, tissue homeostasis and disease^1,2^. For example, sensing of shear flow in blood circulation is essential for the proper development of vasculature and the depletion of rigidity sensors enables cancer neoplastic transformation, i.e. substrate-rigidity-independent growth^3,4^. Mechanosensitive ion channels and cell adhesions, are major mechanosensors that are responsible for converting mechanical cues into biochemical signals in these processes^5^.

A mechanosensing function of focal adhesions is manifest by rigidity-dependent adhesion assembly and turnover, which involves hundreds of different focal adhesion proteins that alter cell behavior during spreading and migration^1^. In addition, focal adhesion mechanosensing regulates pathways such as AKT, MAPK and Hippo-signaling that influence cell fate decisions^1,6,7^. Because adhesions affect cell behavior and fate, the molecular components and dynamics of focal adhesions have been extensively studied. There exists a robust process of adhesion protein recruitment and processing in the finely tuned steps of focal adhesion formation, growth and turnover that correlates with cell state decision making based on cell-microenvironment interactions^8^. There is evidence for force-sensing proteins such as talin and vinculin regulating the functions of focal adhesions as well^2,8–13^. In contrast, although the ion conduction properties of many mechanosensitive ion channels are well-characterized by structural and patch-clamp studies, the mechanisms for converting matrix forces to channel gating are not well understood^14^. This is partly due to the complex nature of ion channel signaling and mechanically-induced calcium influx. For example, mechanosensitive channels are linked to both cell proliferation and apoptosis depending on the context, suggesting that both the distribution and dynamics of the mechanosensitive ion channels are important for understanding the critical mechanosensing processes of the cell.^15,16^

There is ample evidence of a crosstalk between mechanosensitive ion channels and focal adhesions. Local calcium entry is critical for focal adhesion assembly and disassembly based upon the calpain dependence of both processes ^17–21^. In adhesion assembly during cell spreading, rigidity sensing involves force-dependent calpain cleavage of talin, which is needed for normal adhesions to form and for normal cells to grow^20^. In contrast, transformed cells do not sense matrix rigidity and adhesions are significantly smaller even though tumor cells develop higher traction forces^22^. This raises questions of whether or not transformed cells use calcium in adhesion assembly like normal cells. In addition, several studies found that adhesion disassembly depends upon force-dependent calpain cleavage^18,19^. Thus, there are critical roles for calcium entry in normal adhesion dynamics that are altered in transformed (tumor) cells and the analysis of the channels involved can reveal important aspects of the life cycle of focal adhesions.

Several calcium-permeable channels have been implicated in focal adhesions dynamics ^23–27^. One of those, Piezo1, is particularly interesting, as early reports of Piezo1 document an interaction with integrins^28^. Piezo1 has subsequently been identified as a bona-fide mechanosensitive ion channel in mammalian cells^29^. Since then, Piezo1 has been shown to have many roles in important biological processes and there is increasing interest in how it functions^15,29–31^. Piezo1 is widely expressed in somatic cell types and is linked to mechanosensing in stem cell lineage determination, blood pressure regulation, activation of innate immunity and vascular development^4,30,32,33^. Piezo1’s activity has been shown to be related to the integrin-linked actin cytoskeleton ^26,34,35^. When Neuro2A cells are mechanically agitated with micron-sized, collagen IV or Matrigel coated beads, the tension threshold required to trigger Piezo-dependent Ca^2+^ entry is significantly reduced^36^. In addition, integrin signaling pathways are critical for proper function of Piezo1 in the context of endothelial cell responses to flow^37^. Consequently, we want to investigate the possible crosstalk between Piezo1 and focal adhesions.

In this study, we show that in normal cell lines Piezo1 is needed for mature adhesion assembly and it localizes to adhesions as the adhesions mature in a calpain-dependent process, which has been previously observed in the context of talin cleavage^20^. In normal cell lines from many tissues, Piezo1 is enriched at focal adhesions in a force-dependent manner and dissociates after the integrin adhesions disassemble. Its localization correlates with local calcium entry and adhesion disassembly in a calpain-dependent process. Alternatively, in transformed cells, Piezo1 is diffusely distributed and does not affect either adhesion assembly or disassembly and there is low adhesion calcium entry. These studies indicate that Piezo1 normally interacts with adhesions transiently during assembly and then its force-dependent recruitment to mature adhesions catalyzes calpain-dependent adhesion disassembly. These dynamic adhesion interactions are lost in transformed cells and are likely a key neoplastic modification.

## Results

### Piezo1 stably localizes to mature adhesions in HFF cells

To investigate the role of Piezo1 in adhesion formation, we followed the distribution and dynamics of Piezo1 localization during cell spreading in Human Foreskin Fibroblast (HFF) cells using total reflection fluorescence microscopy (TIRFM). Transiently expressed Piezo1 tagged with GFP^38^ or mRuby3 and paxillin-BFP were followed during the cell spreading in HFF cells (Figure 1A top panel, supplementary movie 1). In the initial phase of spreading, Piezo1 did not colocalize with cell adhesions marked by paxillin and it appeared concentrated around the nucleus. After 30-60 minutes when the cells developed mature adhesions, Piezo1 dispersed to the entire spread area of the cell and was significantly enriched in maturing adhesions.

**Figure 1.**
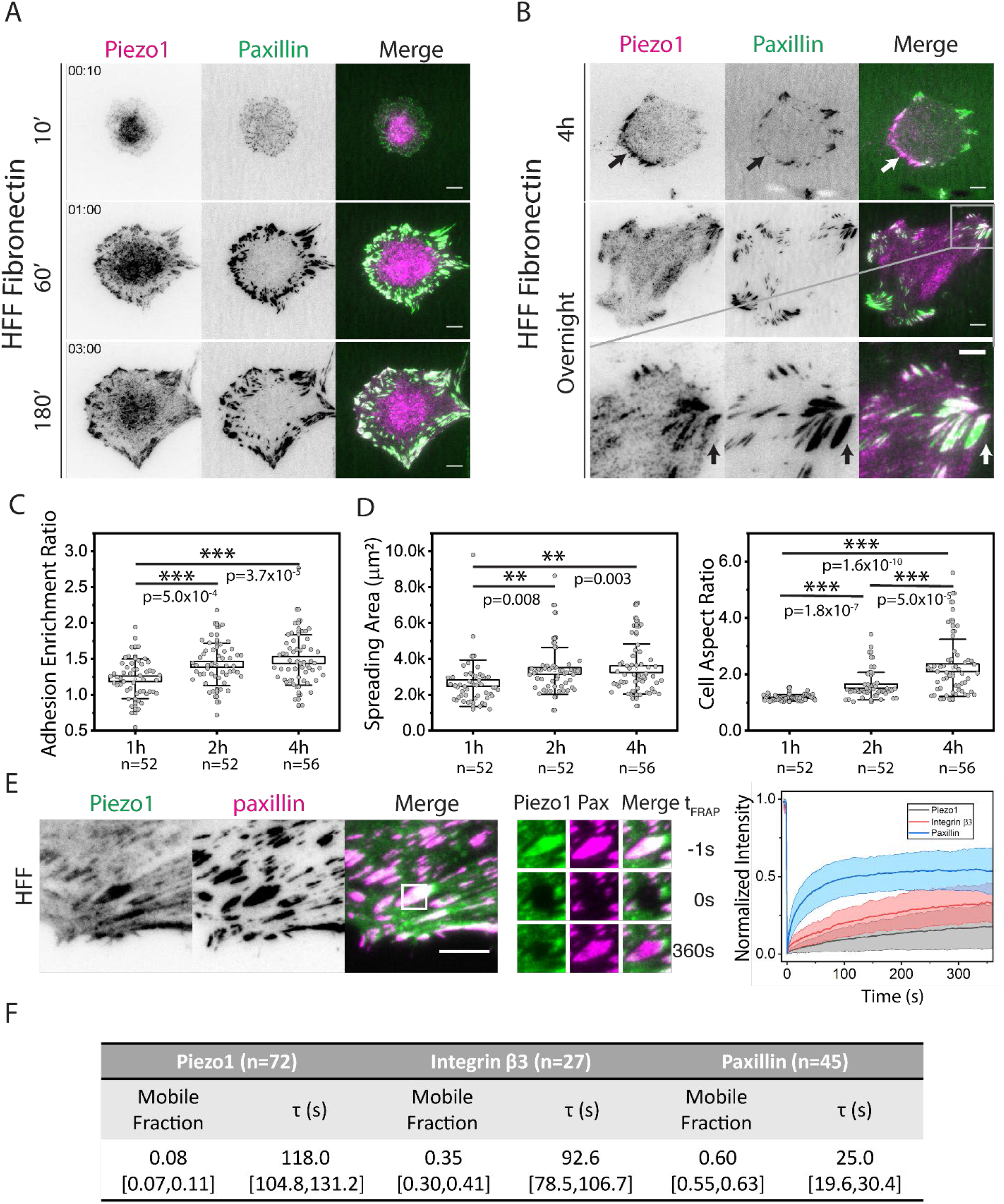
Piezo1 is stably recruited to maturing adhesions on fibronectin surfaces. (A)Time lapse of cell spreading of HFF cells seeded on fibronectin coated glass surface. The cells are transiently expressing Piezo1-mruby3 and paxillin-BFP. Scale Bars denote 10 um. (B) Localization of Piezo1 spreading on fibronectin surface during polarization and in stable condition. Scale Bars denote 10 um in the main figure and 5um in the zoom. (C) Adhesion enrichment of Piezo1 at 1h, 2h and 4h after cell seeding. The adhesion enrichment was defined as the average intesity ratio of Piezo1 in the adhesions and background. (D) Cell spreading area and aspect ratio 1h, 2h and 4h after cell seeding. The significance were calculated using two sample t-test. (E) Left panel: Live TIRF images of Human Foreskin Fibroblast (HFF) cells transiently transfected with Piezo1-GFP and paxillin-mapple on fibronectin coated surfaces. The adhesion region marked by white square was bleached and snapshots of fluorescent recovery for Piezo1 and talin are shown in the zoom in figure on bottom. Scale Bars denote 5 um. Right panel: The FRAP recovery curve of the bleached region (mean ± std). (F) The fitted mobile fraction and recovery rates from the FRAP measurements. More than five independent cells and twenty adhesions were picked for each condition. The bracket after the fitting value denote 95% confidence interval of the fitting parameters. In the Frap experiment, Piezo1 is either co-expressed with Integrin β3 or Paxillin.

Piezo1 localization to adhesions increased over time until the cell began to polarize. Upon cell polarization, Piezo1 prominently localized to the retracting edges (Figure 1B, upper panel). After spreading overnight, HFF cells reached a dynamic steady state and Piezo1 was enriched at mature focal adhesions (Figure 1B, middle panel). In most cells, Piezo1 was recruited to a sub population of the focal adhesions, mostly around the cell periphery at the ends of stress fibers. Quantification of the extent of Piezo1 recuitment to focal adhesions by the ratio of Piezo1 intensity in the adhesion to that in the plasma membrane background showed that the recruitment of Piezo1 to adhesions increased over time as cells spread further and started to polarize (Figure 1 C-D). Piezo1 was often recruited to a portion of individual adhesions as well (Figure 1B lower panel, marked by arrow), implying that the regulation of Piezo1 recruitment happened locally. The localization of endogenous Piezo1 to adhesions in HFF cells was also observed by Piezo1 immunostaining of fixed HFF cells (Figure S1) using two knockdown/knockout validated antibodies^15^. Thus, Piezo1 complexed with mature and retracting adhesions in polarized cells on fibronectin.

Since Piezo1 was previously reported to be diffusive in plasma membranes^26^, we confirmed that the Piezo1 recruitment involved immobilization by checking the lifetime of Piezo1 in the focal adhesions of HFF cells through Fluorescence Recovery After Photobleaching (FRAP). Surprisingly, as shown in Figure 1E, Piezo1 was stable at the adhesions after recruitment, with lifetimes of around 120 s that were considerably longer compared to typical focal adhesion proteins such as paxillin ∼25 s and even slightly longer than integrins such as integrin β3 ∼90 s (Figure 1F). In addition, the mobile fraction of Piezo1 was considerably less than the mobile fraction of paxillin and even integrin β3 (Figure 1F). These data indicated that Piezo1 formed stable interactions with certain focal adhesion components in HFF cells.

### Piezo1 recruitment to adhesions requires myosin II contractility

Since focal adhesion maturation was linked to sustained traction forces and correlated with Piezo1 recruitment during spreading, we hypothesized that Piezo1-adhesion interactions were force dependent. Treatment of Piezo1-GFP expressing HFF cells with 30 uM of Y23762 to inhibit actomyosin contraction, caused rapid loss of Piezo1 from adhesion sites and was followed by the disassembly of focal adhesions as measured by paxillin (Figure 2A, top panel, supplementary movie2). Similar results were observed with 50 μM blebbistatin treatment (Figure S2A). When Y-23762 was washed out, Piezo1 returned rapidly to the reforming adhesions (Figure 2A, top panel). Since Piezo1 recruitment during recovery was much more rapid than the initial recruitment to adhesions during cell spreading (minutes versus hours), we postulated that components remaining at adhesions in the absence of force catalyzed the rapid recovery of adhesion proteins with the development of contractile forces to enable Piezo1 binding. Further, when the contractility of the cell was enhanced by co-expressing the constitutively active version of RhoA along with Piezo1 in HFF cells, the enrichment of Piezo1 at adhesions was strongly promoted (Figure 2A, bottom panel).

**Figure 2.**
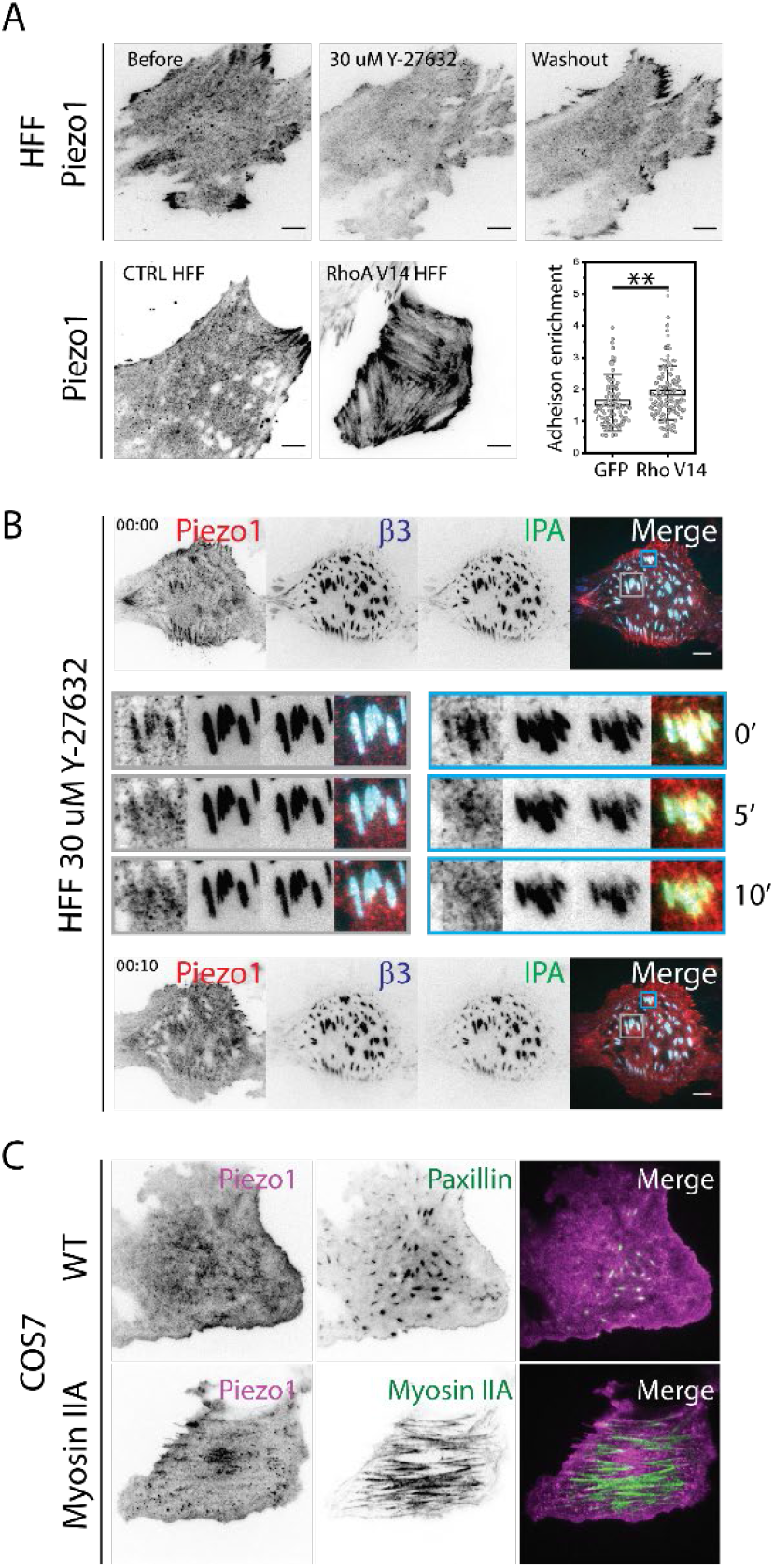
Piezo1 localization in HFF cells is controlled by contractility. (A) Upper panel: Distribution of Piezo1-mRuby3 in HFF cells 15 min after treatment with 30 uM Y-compound and 30 min after washout. Experiments were repeated on > 5 cells with similar results. Lower panel: representative Piezo1 localization and Piezo1 adhesion enrichment quantification in control HFF cells and HFF cells expressing constitutive-active RhoA V14 mutant. Scale Bars denote 10 μm. (B) Time-lapse image of HFF cells transfected with Piezo1-mruby3, integrin β3-Emerald and IPA-GFP with the addition of 30 μM Y-27632. Experiments were repeated on > 3 cells with similar results. (C)Fluorescence image of Piezo1-GFP in COS7 cells with/without overexpressing myosin IIA-mCherry.

To check whether the reduction of traction forces alone could disrupt Piezo1’s adhesion recruitment without disruption of adhesion proteins such as talin and integrins, we tested the relationship between Piezo1 localization and myosin II contractility in intact adhesions. Previous reports showed that inhibition of the tyrosine phosphatase, PTP-PEST, as well as the expression of virulence bacterial protein IpaA, inhibited adhesion disassembly upon addition of Y-23762 ^39,40^. In cells expressing GFP-labelled IpaA peptide, addition of 30 μM Y-23762 caused cells to lose contractility while retaining adhesion structures (Figure 2B, Supplemental movie 3). In this case, Piezo1-mRuby3 diffused out from adhesions within 10 min after Y-23762 addition while the integrin β3-BFP remained intact. A similar effect was observed when 2.5 μM of PTP-PEST inhibitor was added in combination with 30 μM Y-23762 (Figure S2B). Thus, the localization of Piezo1 to focal adhesions depended on continued contraction forces possibly through the stretching of adhesion proteins, and was independent of paxillin or integrin β3 localization to adhesions.

To further establish the role of myosin II-based contractility in Piezo1 recruitment, we checked the distribution of Piezo1 in cells that had low traction forces such as COS-7 cells lacking myosin IIA expression^41^. In COS-7 cells transiently expressing Piezo1-GFP and paxillin-mapple, Piezo1 was diffusive in the membrane with no apparent localization to adhesions (Figure 2C, top panel) and the cell adopted a highly-spread disc morphology. When contractility was restored in COS-7 cells by overexpressing myosin IIA, cells became polarized with myosin IIA forming stress-fiber-like cables (Figure 2C, bottom panel) and Piezo1 was recruited to the adhesions at the ends of the myosin cables. This further reinforced the hypothesis that Piezo1 recruitment to adhesions required continued force on the adhesion complex.

### Piezo1 recruitment triggers (calpain dependent) integrin β3 adhesion turnover

Since Piezo1 was recruited to only a fraction of focal adhesions, we proceeded to check whether different integrin subtypes played a role in the regulation of Piezo1 localization. As shown in Figure 3A, we transiently expressed Piezo1-1591-mRuby3 with integrin β3-mEmerald and analyzed the colocalization of Piezo1 with β3 adhesions during cycles of adhesion assembly and disassembly. Interestingly, the recruitment of Piezo1 to β3-enriched adhesions correlated with the disassembly and turnover of the β3 adhesions. Figure 3A shows the timelapse views of two such turnover events on fibronectin surfaces, one during spreading where the cell was protruding and the other in migrating cells where the adhesion was retracting. In both cases, integrin β3-based adhesions formed without apparent Piezo1 co-localization. Over time, Piezo1 was recruited to the adhesions from the periphery of the adhesions and gradually occupied the whole adhesion. Piezo 1 recruitment appeared to initiate a decrease in integrin β3 fluorescence intensity within minutes (Figure 3A and supplementary movie 4-5). After integrin β3 reached background levels, the Piezo1 fluorescence often persisted in adhesions for a few minutes before dissociation (Figure 3A, Left panel, arrows; Figure S3A gives a detailed profile), suggesting that some adhesion components remained that supported Piezo1 localization after integrin β3 dissociated from the adhesion.

**Figure 3.**
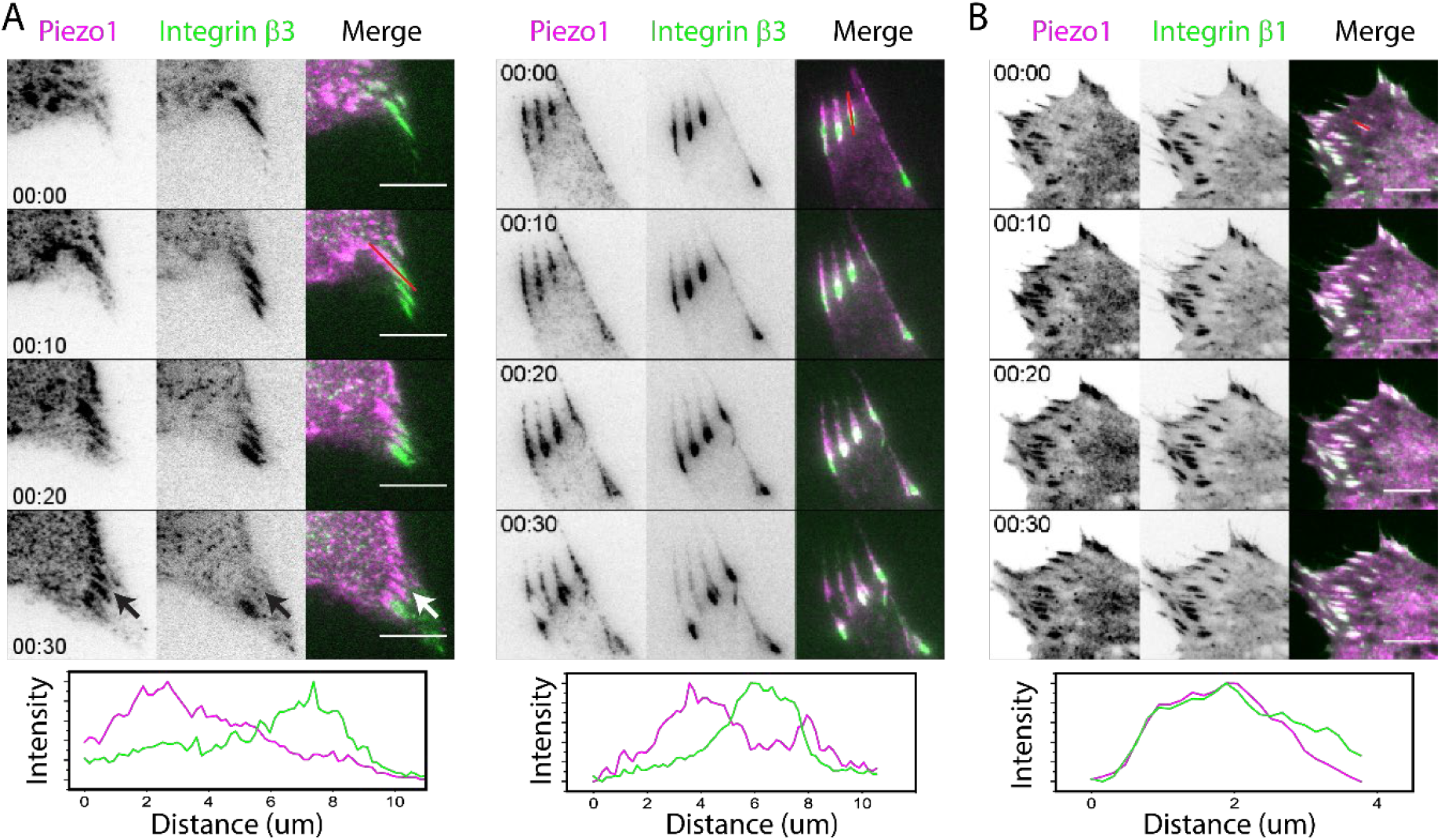
Piezo1 recruitment of adhesions precede integrin β3 disassembly. (A) Kymographs of HFF cells co-transfected with Piezo1-mruby3 and mEmerald-Integrin β3 spreading on fibronectin surfaces. Left panel: A spreading cell one hour after seeding. Right panel: A retracting cell after overnight seeding. (B) Kymographs of HFF cells transfected with Piezo1-mruby3 and mEmerald-Integrin β1 spreading on fibronectin surfaces overnight. The normalized intensity profile of adhesions highlighted by the red line were shown below the figure.

In contrast, the anticorrelated behavior was not observed between Piezo1 and β1 integrins in adhesions. Piezo1 recruitment at integrin β3 and β1 enriched adhesions was more stable, and they often co-localized for tens of minutes (Figure S3B). In an interesting example (Figure S3B), Piezo1 recruitment correlated with the rapid dissociation of β3 integrins and enrichment of β1 integrins. Piezo1 recruitment persisted in adhesions even after dissociation of β3 integrins (Fig. S3B, arrows). These observations indicated that Piezo1 recruitment was associated with adhesion turnover that was regulated locally in an integrin β3 dependent manner.

Considering Piezo1’s function as a mechanosensitive Ca^2+^ channel, adhesion turnover was likely caused by the calpain-dependent cleavage pathway. Indeed, when we blocked calpain activity using the inhibitor ALLN, adhesion turnover from the back of the sliding adhesions was inhibited, causing significant elongation of the adhesions (Figure S3C, left panel, Supplementary Movie 6). However, the speed of adhesion extension was not altered (Supplementary Movie 6). Piezo1 remained in adhesions following ALLN addition in HFF (Figure S3C). Thus, normal dynamics of β3 adhesions depended upon calpain activity that was likely regulated by local Ca^2+^ entry induced by Piezo1’s channel activity.

Based upon these findings we hypothesized that Piezo1 recruitment was upstream of calpain in regulating adhesion turnover. In earlier studies of cells on fibronectin, integrin β3 adhesions formed initially, and then the adhesions switched to integrin β1. The greater stability of β1-containing adhesions with Piezo1 recruitment could potentially provide a mechanistic explanation for such switching events^42^.

### Piezo1 localization to adhesions is lost in transformed cells

To check whether Piezo1 localization to adhesions was cell-type specific, we expressed Piezo1 in a variety of normal and transformed cell lines (Figure 4A, Figure S4A). In all of the normal cell lines of fibroblastic (HFF, MCF-10A), epithelial (HUMEC) or endothelial origins (HUVEC), Piezo1 localized to a significant subset of the focal adhesions on fibronectin visualized by paxillin (Figure 4A, Figure S4). In the case of 12 different tumor cell lines from mesenchymal or epithelial tissues (fibrosarcoma, melanoma, breast and lung cancers, Figure 4D), there was no significant colocalization of Piezo1 with paxillin adhesions. In the HT1080 (fibrosarcoma) and MDA-MB-231 (breast cancer) cells, FRAP measurements of Piezo1 diffusion showed that Piezo1 had a significantly higher mobile fraction than in HFF cells (Figure 4B). The difference in Piezo1 immobilization was not due to different levels of Piezo1 expression, since Western blots showed significant levels of Piezo1 in most of these cell lines (Figure 1D, Figure S4B). Thus, the colocalization of Piezo1 with adhesions was found only in normal cells. In all of the tumor cells tested, Piezo1 was not colocalizing with adhesions and was rapidly diffusing.

**Figure 4.**
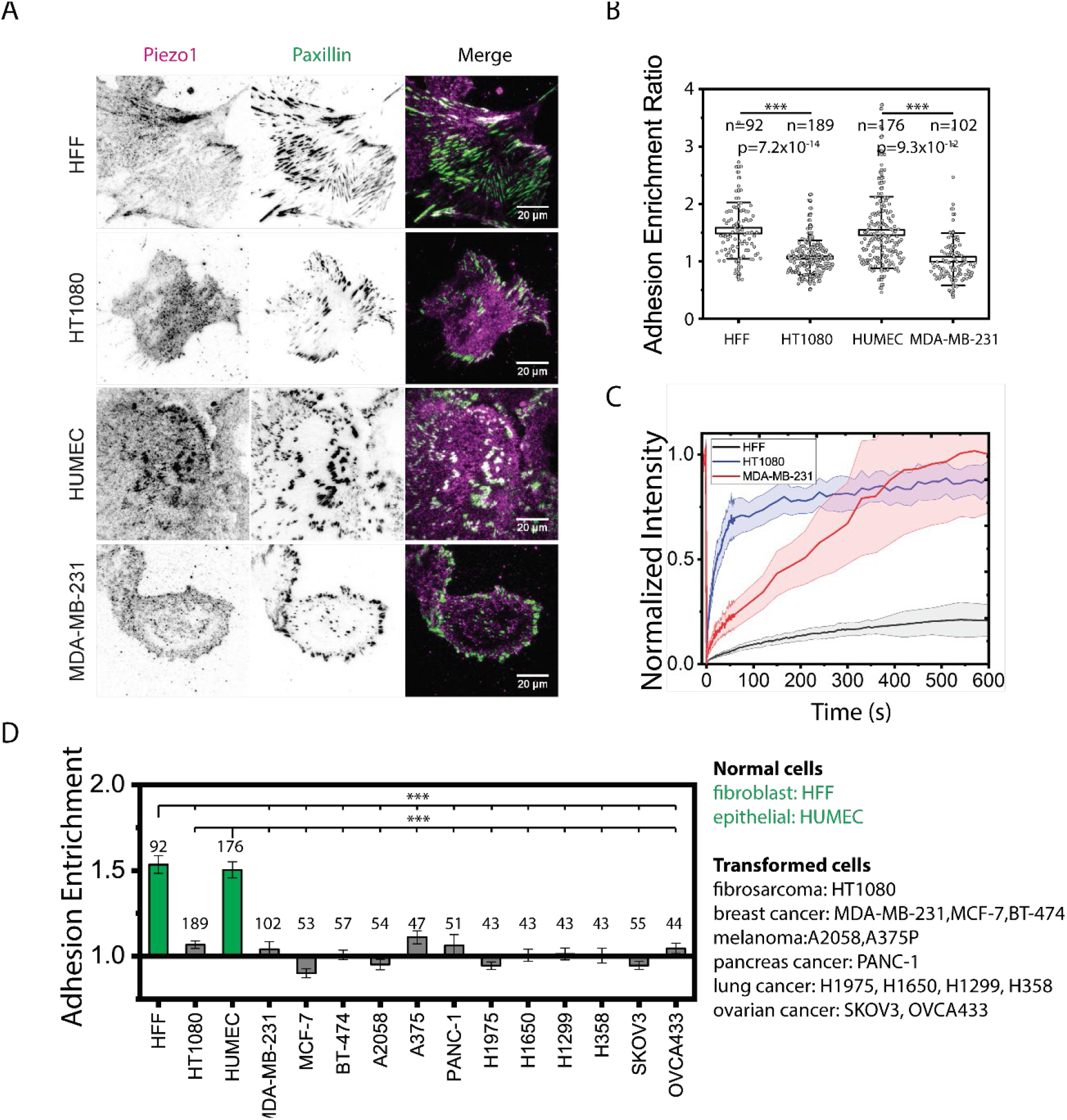
Piezo1 localizes to the adhesions of non-transformed cells but is diffusive in transformed cells. (A) Immunostaining of Piezo1 localization in paired normal and transformed cell lines with similar orgin. (B) Quantification of Piezo1 adhesion enrichment for cells in Figure (A). p-values were calculated using two-sample t-test. (C) FRAP recovery curve of Piezo1 in HFF cells and transformed HT1080 and MDA-MB-231 cells. The error bar denotes 95% confidence interval of the mean. (D) Quantification of adhesion entrichment of Piezo1 across various normal and transformed cancer cell lines. Error bars denote standard error of the mean. There were statistically significant differences (p-value < 0.001) between HFF/HUMEC and all the cancer cells.

Our previous studies showed that TPM 2.1 levels were low in many transformed cells^22^. Knocking down TPM 2.1 in HFF cells caused the cells to become transformed with smaller adhesions and the cells were not able to differentiate between soft and rigid surfaces. In contrast, restoring TPM 2.1 expression in MDA-MB-231 cells caused normal behavior with bigger adhesions and the cells died on soft agar^22^. Based on this, we checked if toggling between the non-transformed and transformed states in HFF and MDA-MB-231 cells altered the localization of Piezo1 in these cell lines. When we knocked down TPM 2.1 in HFF cells, the recruitment of Piezo1 to adhesions was significantly weakened compared with control cells or control siRNA-treated cells (Figure S5 A). In contrast, when we transiently expressed TPM 2.1-YFP in MDA-MB-231 cells, a fraction of cells formed stress fibers and Piezo1 was recruited to adhesions at the end of stress fibers (Figure S5 B). These data indicated that there was a correlation between normal cell behavior and Piezo1 adhesion localization and we recently showed that transformation resulted from the loss of rigidity sensing, which was observed in the majority tumor cells^22^. Thus, the change in cell state from normal to transformed state diminished Piezo1 binding to adhesions.

### Piezo1 is needed for adhesion maturation and polarity in normal but not in transformed cells

As the pattern of Piezo1 adhesion recruitment differs in normal and transformed cell lines, we checked if Piezo1 regulated focal adhesion morphology differently in normal and transformed cell lines. To minimize the possibility of cell-type variance, we used normal primary cells and transformed cancer cells of matching origin (Human foreskin fibroblast with HT1080 human fibrosarcoma). As shown in Figure 5A, Piezo1 knock-down HFF cells spread to smaller areas, had a lower aspect ratio and had smaller adhesions, while in the transformed HT1080 cells, knocking down of Piezo1 caused a significant increase in cell spread area but had no effect on aspect ratio or adhesion size (Figure 1 A-C).

**Figure 5.**
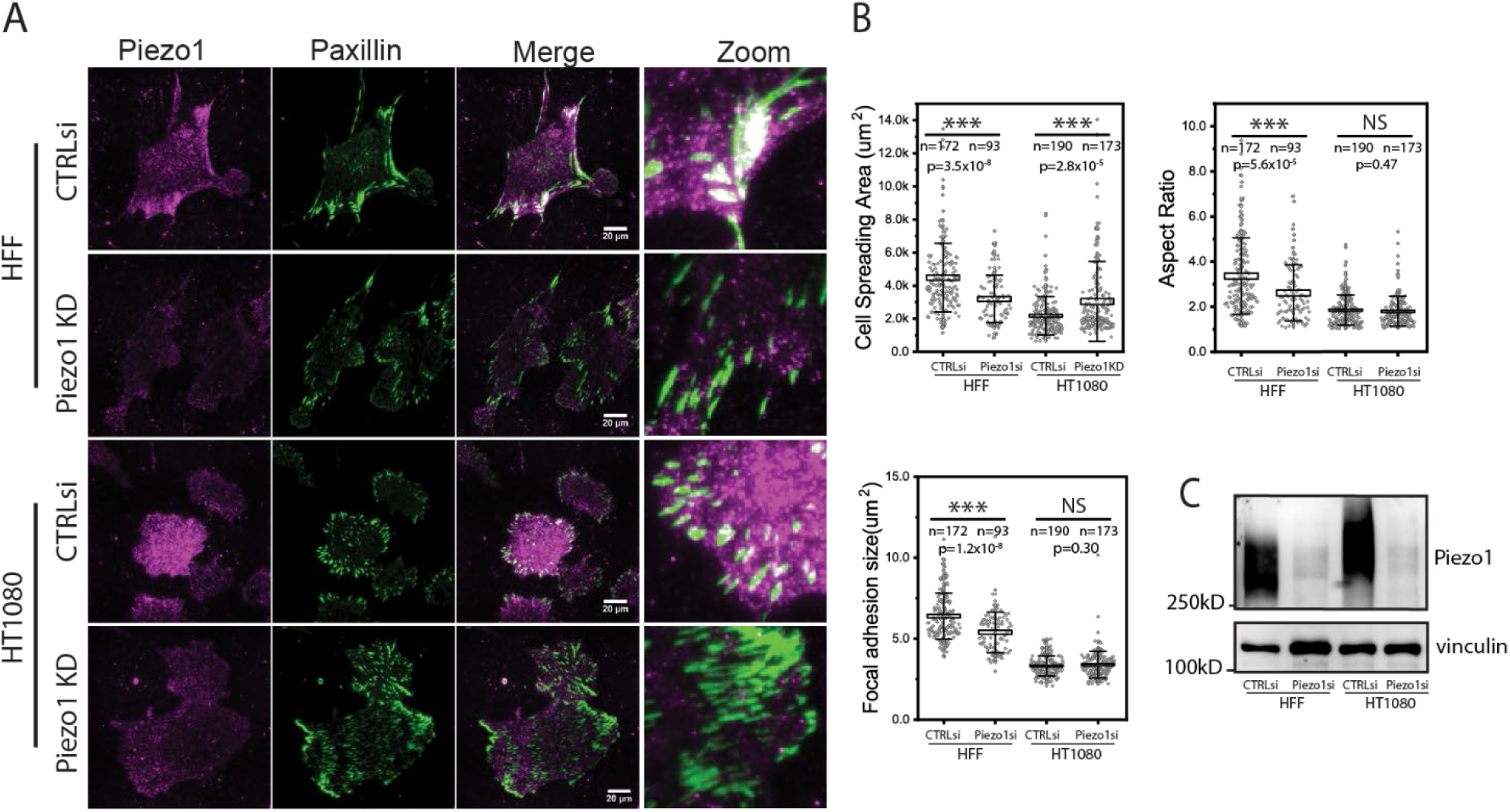
Piezo1 localization affects adhesion maturation and correlates with transformation states of cells. (A) TIRF image of fixed cell staining of non-transformed HFF cell line and transformed fibrosarcoma HT1080 with/without Piezo1 knock down after overnight spreading on fibronectin surfaces. Scale Bars denote 20 um. (B) Box plots of the cell spreading areas, cell aspect ratio and mean adhesion sizes of the cells in figure A. Box and error bar denotes the s.e.m and s.t.d of the measurements. The significance value were calculated using two sample t-tests. (C) Western blots of Piezo1 level in control siRNA and piezo1 knock-down cells.

This indicated that Piezo1 was important for adhesion maturation and cell polarization in normal cells but had little effect on adhesion formation in transformed cells, while affecting spreading area. Previous studies documented that transformed cell lines including cancer cells generally had smaller and less mature adhesions compared with non-transformed cells^22,43^. This was despite the fact that the majority of adhesion components were present at similar levels in both normal and transformed cells^44,45^. Thus, Piezo1 recruitment to adhesions in normal cells appeared to be involved in important steps in the adhesion maturation process whereas in transformed cells, Piezo1 depletion caused no change in adhesion morphology^46^.

### Piezo1’s adhesion recruitment leads to local calcium transients around adhesions in normal but not in transformed cells

Next we checked the possible physiological significance of the Piezo1 adhesion recruitment. Since Piezo1 functions as a mechanosensitive ion channel, the recruitment of Piezo1 to adhesions may have altered the calcium entry pattern of the cells. Recent studies by Ellefsen et. al, showed that Piezo1-dependent calcium spikes were localized to the cell periphery near focal adhesions^26^. This implied that calcium spikes occurred near where there were concentrations of Piezo1 at adhesions. Because Piezo1 was not concentrated in adhesions of transformed cells such as MDA-MB-231 and HEK293T, there was a question of whether or not similar calcium spikes would occur near adhesions in transformed cells.

To determine if calcium entry near adhesions in fibroblasts also occurred in tumor cells, we loaded HFF and MDA-MB-231 cells with the cell-permeable calcium indicator Cal-520 and imaged transient calcium “puffs” on fibronectin surfaces. Similar to previous observations in HFF cells^26^, many local calcium entry events were detected in areas of high contractility at the cell edge and surrounding adhesions (Figure S6). Interestingly, although the transformed MDA-MB-231 cells generated higher local traction forces^22^ and were killed by calcium-dependent apoptosis after stretch^16^, they had very few calcium spikes on fibronectin surfaces (Figure S6). This indicated that the calcium influx in cancer cells was abnormal and was consistent with the lack of Piezo1 within adhesions after transformation.

Although the measurements with Cal-520 calcium sensor provided interesting insights into the differences in spatial calcium signaling between normal and transformed cells, there remained several specific questions about whether these calcium fluctuations were localized to sites of adhesion. The diffusive nature of all Ca^2+^ dyes reduces the signal-to-noise ratio for calcium influx at sub-cellular locations, providing only limited spatio-temporal information on local Ca^2+^ changes. In addition, under imaging conditions sensitive to subcellular calcium events, Cal-520 had significant photo toxicity that caused rapid increases in intracellular calcium levels limiting the observation window. To better quantify and understand the details of calcium entry at the adhesions, we generated a novel focal adhesion targeted calcium sensor, in which one of the latest most sensitive recombinant calcium sensors, jGCaMP7s, was linked to a paxillin mScarlet-I construct^68^. This improved the signal-to-noise ratio specifically at the adhesions, and the ratio between the calcium sensor and the mscarlet-I fluorescence enabled normalization of the calcium levels based on the local sensor concentration.

When the paxillin calcium sensor was transfected in HFF, HT1080 and MDA-MB-231 cells, it localized robustly to apparent adhesions. After overnight spreading, the intensity ratio of jGCaMP7s and mScarlet-I in the focal adhesions of HFF cells showed frequent calcium spikes (Figure 6 A, supplementary movie 7). The calcium fluctuations were greatly reduced in Piezo1 knock down cells, indicating that Piezo1 was required for the calcium fluctuations around the adhesions(Figure 6 A, supplementary movie 8). Interestingly, in the transformed HT1080 and MDA-MB-231 cells the calcium spikes at the adhesions were significantly weaker, often there was no detectable calcium signal during a 1-hour observation window(Figure 6 A, supplementary movie 9-10). In the transformed cells, matrix adhesions still formed and disassembled, suggesting that calcium entry around adhesions was not required for adhesion turnover in the transformed cells. These results indicated that Piezo1 recruitment to adhesions in HFF cells enabled calcium spikes that could in turn induce calpain-dependent adhesion turnover; but in transformed cells, adhesion dynamics and cellular calcium fluctuations were de-coupled.

**Figure 6.**
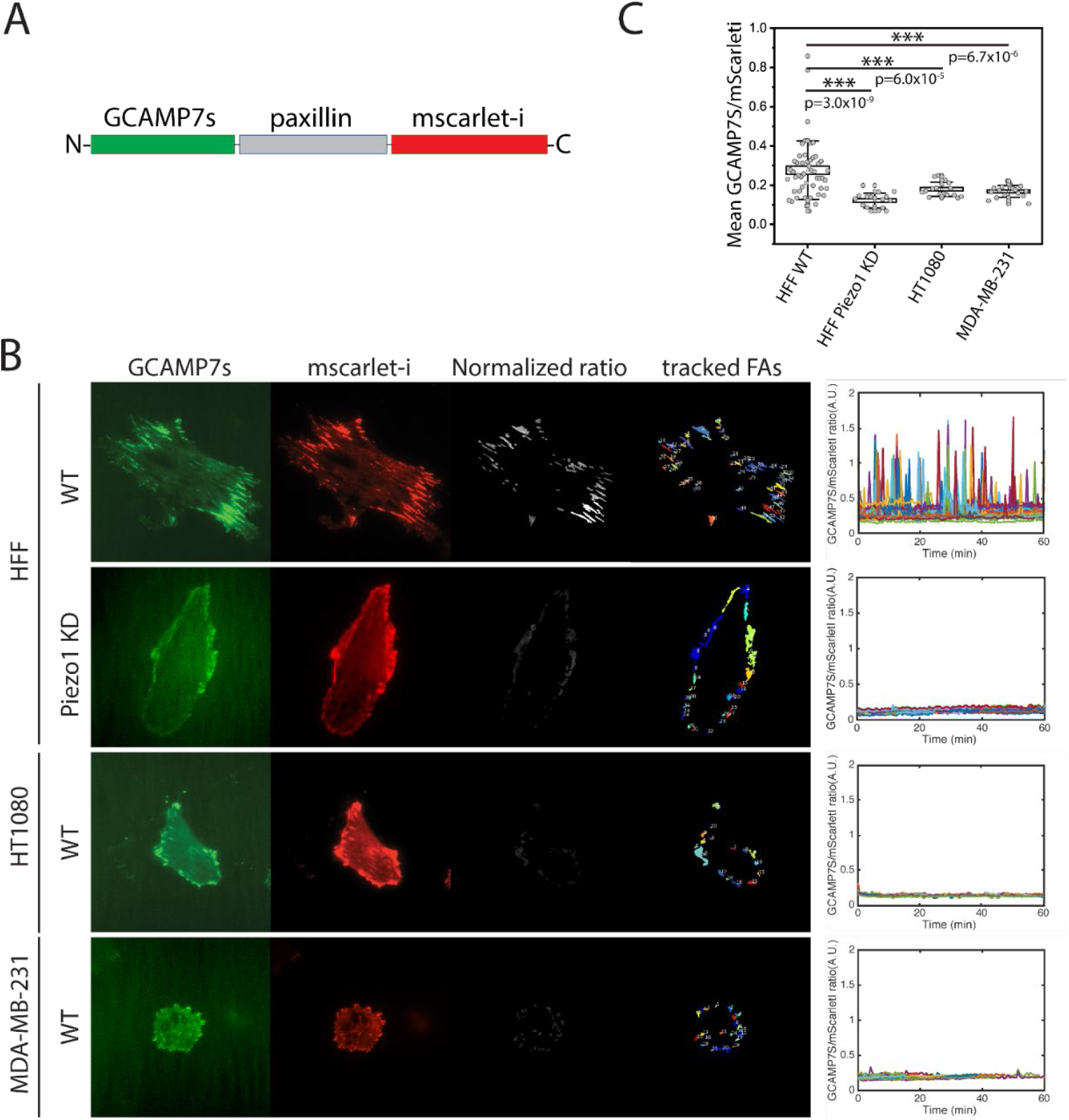
Normal and transformed cells have distinct adhesion calcium signals. (A) Domain Illustration of the novel adhesion targeting calcium sensor based on jGCaMP7s from Janelia. (B) Left: TIRF image montages of representative HFF cells (with/without Piezo1 knock down), HT1080 and MDA-MB-231 cells transfected with paxillin calcium sensor. The cells were seed on fibronectin surfaces overnight before imaging. Normalized ratio represents the ratio image between jGCaMP7s and mScarlet-I, which is an measure of local Ca^2+^ concentrations. Right: calcium level fluctuations over time in the tracked adhesions of the cell presented in the left panel, each line in the image denotes a single adhesion. The interval of measurement is 30 s. (C) Quantification of average calcium ratio in tracked adhesions over 60 min for different cell types.

### The central linker domain (1418-1656) of Piezo1 is required for Piezo1 adhesion recruitment and alters cell morphology of non-transformed cells

We proceeded to investigate potential molecular mechanisms of Piezo1 adhesion recruitment. So far, no binding partners of Piezo1 at focal adhesions have been reported in the literature. The long lifetime of Piezo1 at the adhesions and the dependence of Piezo1 recruitment on actomyosin contractility led us to hypothesize that Piezo1 bound to adhesion-linked components of the actin cytoskeleton stretched by cellular contractility. Upon examining the 3D architecture of Piezo1^47^, we identified a long intracellular linker as a potential interaction site between Piezo1 and the actin cytoskeleton.

This long intracellular linker region was largely unresolved in the Cryo-EM structures and connected the beam to the clasp and via another unresolved loop to Piezo1 repeat B^47,48^. Previous biophysical studies showed that deletions at the proximal portion of the loop influenced the biophysical properties of the channel via effects on the latch domain^49^. However the distal end was not explored in those studies.

To check the potential function of the linker region of Piezo1, we first made a mNeongreen construct of the larger region of human Piezo1 (residue 1418-1656;mNeonlinker) that contained the unstructured loop region, the clasp and a second disordered loop previously shown to be able to house fluorescent tags^38^. We next checked for its localization and interaction with Piezo1. As shown in Figure 7A, the mNeonlinker construct expressed in HFF cells appeared to be diffusely distributed with no apparent concentrations. However, Piezo1 recruitment to adhesions was reduced as was adhesion size with overexpression of the mNeonlinker (Figure 7A). This is consistent with the hypothesis that at high expression levels the mNeonlinker domain acted as a dominant negative competitor and displaced Piezo1 from adhesions. When we checked the morphology of the cells, HFF cells had significantly lower cell spread area and aspect ratio with the mNeonlinker over-expression, which resembled the morphology change of HFF cells after Piezo1 knock-down. These results were consistent with the hypothesis that the recruitment of Piezo1 to focal adhesions in non-transformed cells activated local calcium fluctuations in surrounding adhesions, and the calcium signal led to adhesion maturation and cell polarization. In the case of transformed cells where Piezo1 did not localize to adhesions, expression of mNeonlinker should have caused minimal effect on cell morphology. Indeed, expression of mNeonlinker in HT1080 cells caused no change in morphology, cell spread area or aspect ratio compared with controls (Figure 7A-C).

**Figure 7.**
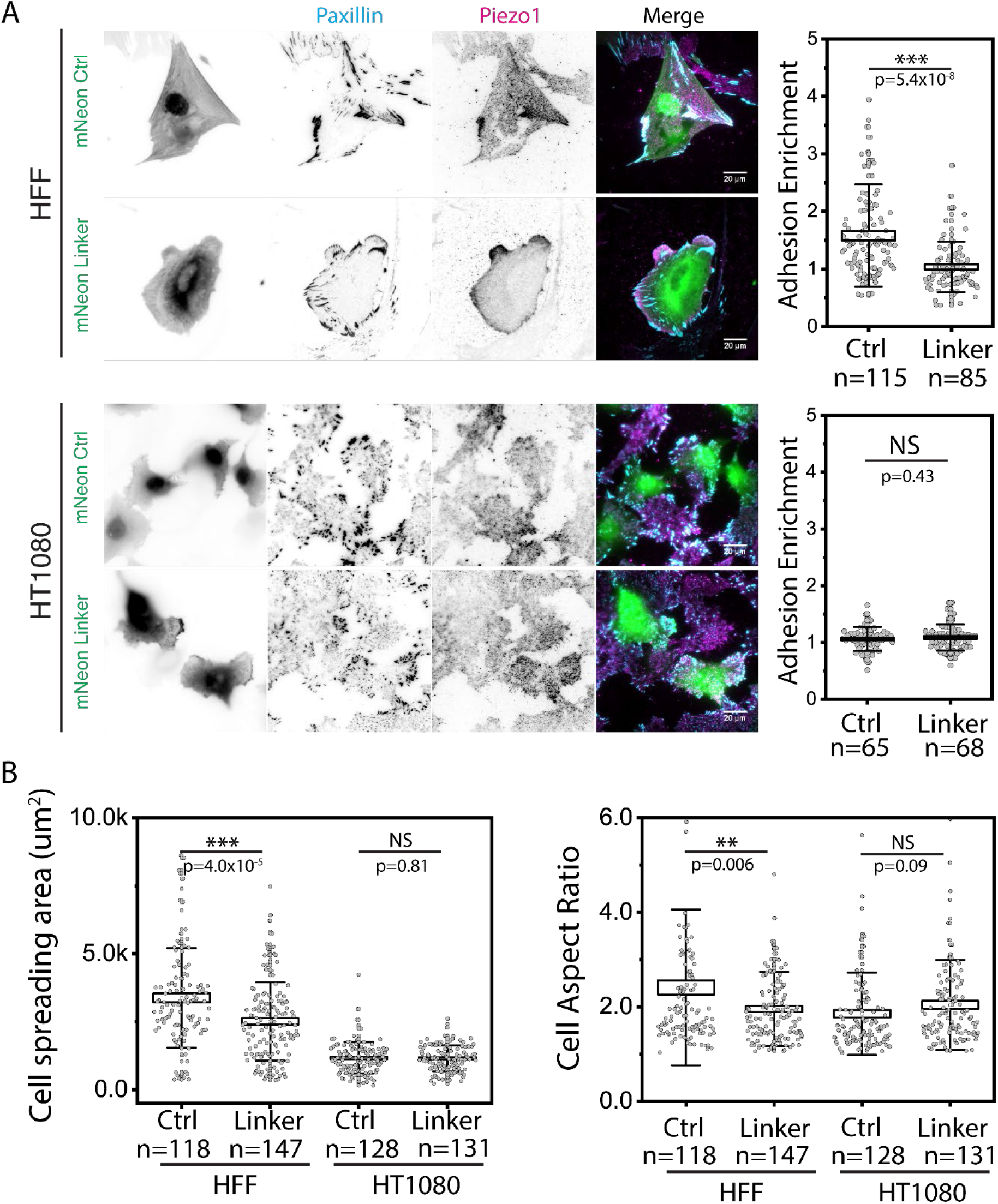
The linker domain of Piezo1 is linked to adhesion recruitment and alters cell morphology in HFF cells. (A) Immunostaining of HFF and HT1080 cells transfected with mNeongreen control vector or mNeongreen labeled Piezo1 linker construct. The cells were seeded on fibronectin surface overnight and stained with paxillin and Piezo1. (B) Box plot of cell spreading area and cell aspect ratio of positively transfected cells. Box and error bar present s.e.m and s.t.d respectively. Statistical significance were calculated using two sample t-test.

To further check if this linker region of Piezo1 indeed harboured focal adhesion binding sites, we generated several Piezo1 mutants with the linker regions deleted (short: Piezo1 Δ1443-1556and long: Piezo1 Δ1443-1473). The Δ1443-1473 deletion removed the loop region and the longer Δ1443-1556 mutant removed much of the clasp domain. When we expressed these deletion mutants in FAK -/- MEF cells that had prominment recruitment of WT Piezo1 to adhesions, we observed normal Piezo1 adhesion recruitment of Piezo1 Δ1443-1473 mutant and diffusive Piezo1 localization of the Δ1443-1556 mutant (Figure S7 and supplementary movie11-12). Electrophysiological characterization in Piezo1 KO HEK293T cells showed that the Δ1443-1473 mutant had normal stretch induced channel activity while the Δ1443-1556 mutant was not functional.

Taking these data together, it appeared that the linker domain(residues 1418-1656) and most likely the clasp region of Piezo1 harbored a binding site for focal adhesion components that recruited Piezo1 to focal adhesions in a force-dependent manner in non-transformed cells at the membrane-cytoplasm interface.

## Discussion

In this study, we show that Piezo1 has important functions in focal adhesion dynamics in normal cells but not in transformed cells. Normal fibroblasts, epithelial and endothelial cells form focal adhesions on fibronectin where Piezo1 concentrates in a force-dependent process. The lifetime of Piezo1 in the adhesions is similar to that of integrins and longer than typical adhesion proteins, which implies that it complexes stably with adhesion bound proteins. Sustained contraction forces are needed for piezo1 localization to adhesions; and when contraction is inhibited, it could leave adhesions before the dissociation of adhesion components such as integrin and talin. Piezo1 localization to focal adhesions leads to the dissociation of integrin β3 containing adhesions but not integrin β1 adhesions on fibronectin. This localization is also lost in transformed cells, including virus-transformed somatic cells (e.g. HEK293T) as well as tumor cell lines from different tissues. When somatic cells are transformed such as in the case of TPM 2.1 knock down, the recruitment of Piezo1 to adhesions is dramatically reduced similar to the transformed cancer lines tested. Calcium levels in the vicinity of adhesions are higher and have many more local fluctuations in normal cells compared with transformed cells. Binding to adhesions depends upon an unstructured cytoplasmic domain of Piezo1 (aa 1418-1656). These findings indicate that Piezo1 has a number of complex interactions, which depend upon contractility, the transformed state, as well as specific integrin-matrix interactions. Piezo1 appears to have multiple context-dependent functions in normal and transformed cells that need to be analyzed temporally and spatially in a defined mechanical environment.

Piezo1 was first identified as an integrin-interacting protein which had a role in adhesion formation^28^. Our results reinforce those findings and further indicate that Piezo1 binds in a complex with integrins but not with adhesion proteins such as talin or paxillin, which have much more rapid recovery rates after photobleaching. Further, inhibition of PTPN12 activity or expression of the *Shigella* IpaA effector stabilizes adhesion proteins even after inhibition of myosin but does not prevent Piezo1 dissociation. Piezo1’s rapid dissociation upon myosin inhibition is rapidly reversed upon washout of the myosin inhibitor. The rate of return to the adhesions is much more rapid than during initial adhesion formation, indicating that the recruitment of Piezo1 at adhesions depends on a force-sensitive protein or activation of a force-dependent enzyme that is retained at adhesion sites during myosin inhibition. Overexpression of a linker region of human Piezo1 acts as a dominate negative, displacing Piezo1 from adhesions, implying that this linker region of Piezo1 harbours the interacting site for adhesion binding. The lack of adhesion maturation after depletion of Piezo1 strongly supports the hypothesis that Piezo1 has an important role in the force-dependent maturation of adhesions on rigid surfaces, and this does not occur in transformed cells.

Since the discovery of Piezo1’s role as a mechanosensitive ion channel in eukaryotic cells, the regulation mechanism of Piezo1’s function in the various physiological activities of cells has been under heavy investigation^50,51^. The classical model of Piezo1’s physiological function assumes that it is diffusive in the plasma membrane and is activated locally through membrane bilayer tension or curvature changes that occur through cytoskeletal-dependent mechanical events^26,34,38,47,52^. Several recent studies have clearly shown that the cystoskeletal environment influences Piezo1 function^38^, including the observation that Piezo1 localizes to the adhesions of some glioblastoma cell lines, which facilitates their growth and invasion^27^. The unusual nature of these glioblastoma lines indicates that they may be more like normal cells than transformed cells (Yao et al unpublished results). In our study, we systematically examine Piezo1’s spatial distribution across many cell types from different tissues of origin and discover that Piezo1 adhesion localization is much more prominent in non-transformed normal cells compared with transformed cells such as most tumor cell lines. Binding to the adhesions in normal cells catalyzes adhesion maturation and disassembly that correlates with cell growth.

Interestingly, in contrast to the reduced calcium transients that we observed in resting transformed cells, we recently reported that mechanical perturbations, such as stretching or ultrasound treatment, cause Piezo1-dependent increases in cytoplasmic Ca^2+^ levels and subsequent apoptosis of transformed but not normal cells^53^. These observations are difficult to explain in terms of fluctuations in either mechanical forces on adhesions or membrane tension, especially since fewer Ca^2+^ fluctuation events are observed in transformed cells whereas Ca^2+^ flickering events near adhesion sites are common in normal cells. With regard to membrane tension, typical measurements of membrane tension in cells report values that are 2-3 orders of magnitude smaller than those needed to activate Piezo1 in vitro^54,55^. Certainly, the pattern of Ca^2+^ activation in the HFF cells in the vicinity of adhesions is consistent with previous results^46,56,57^, but it is not consistent with the simplistic model of very high membrane tension for the whole cell. Rather, there appear to be local activation events that involve elements from the adhesions that may alter local membrane tension/curvature or the association of membrane/cytoskeletal proteins that can modulate the mechanical gating of Piezo1 activity^52,58,59^. Although Piezo1 appears in many different mechanical functions, the activation mechanism of Piezo1 is complex and both contributions from membrane tension and interaction with cytoskeletal proteins may modulate its activity^35,58^.

We recently showed that transformed cancer cells are depleted of the rigidity sensing, local contraction units^22,43^. There is a conserved set of cytoskeletal proteins that are able to assemble into local rigidity sensing contractile structures in normal cells. After the depletion of any one of several cytoskeletal proteins critical for rigidity sensing and contractile pair assembly, the cell will be transformed and will no longer sense substrate rigidity. Transformation also involves changes in adhesion morphology and composition^22^. The differences in the adhesion components in normal and transformed cells that contribute to the adhesion phenotype are still an open question but the mRNA levels of over 700 proteins are altered upon transformation by tropomyosin 2.1 depletion^22^. An important function of the rigidity sensors is in the maturation of focal adhesions that contain Piezo1 and without Piezo1 the mature adhesions do not form. This further fits with the absence of any differences between transformed cell adhesions with and without Piezo1. Thus, we suggest that normally both the maturation and the disassembly of focal adhesions is catalyzed by Piezo1. In addition, whether the differences in Piezo1 recruitment in the transformed and somatic cell lines are caused by changes in the molecular interactions or tension states of these cells remains an open question and awaits further investigation.

Our model of the function of Piezo 1 in Figure 8 emphasizes the catalytic role of Ca^2+^ entry in the life cycle of the adhesions in normal cells. Initially, the nascent adhesions seed actin filament formation, myosin bipolar filaments pull on the actin in the rigidity-sensing contractions, and there is a calcium-dependent cleavage of talin1 that is needed for growth^8,20^. If the matrix is rigid, then adhesions mature through enzymatic modifications that trigger the maturation process. Components in the adhesions are stretched by traction forces and then can bind Piezo1 which permits Ca^2+^ entry in a positive feedback cycle for the further maturation of the adhesion. However, when the concentration of Piezo1 reaches a threshold, there is excessive Ca^2+^ entry and recruitment of calpain that now hydrolyzes components of the adhesion, causing the disassembly of the adhesion. The maturation process requires the rigidity sensing complex and will not occur in transformed (cancer) cells. This model indicates that covalent modification of adhesion components during maturation as well as traction forces on those components are necessary for Piezo1 binding. Further understanding of Piezo1 functions requires dynamic studies of Piezo1 at the appropriate time and location during cell spreading or migration.

**Figure 8.**
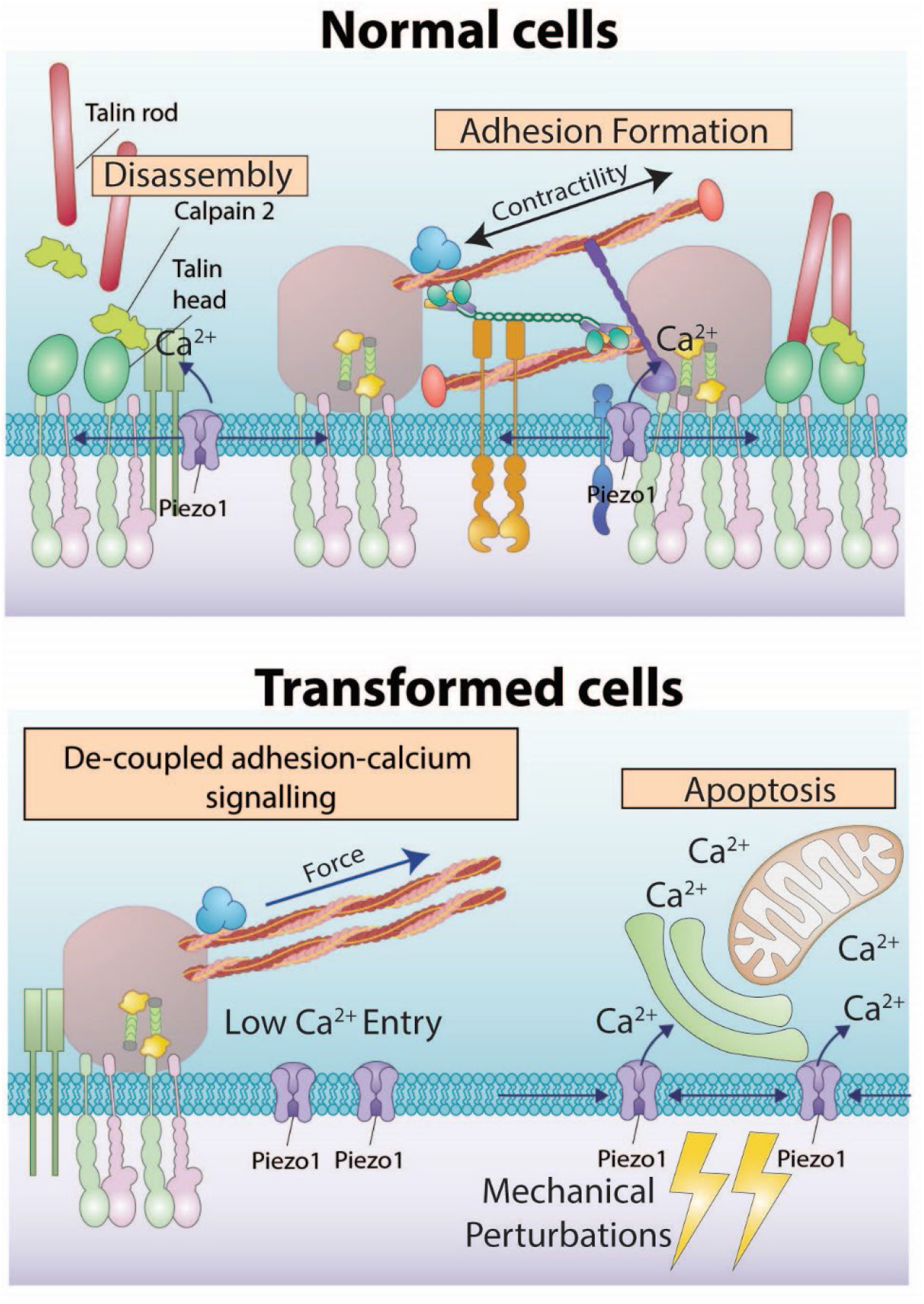
Model of the Piezo1’s regulation in normal and transformed cells. (Top Panel) In normal cells, Piezo1 is recruited to maturing focal adhesions in a force dependent manner, where it regulates both the growth and turnover of focal adhesions. (Bottom Panel) In transformed cells Piezo1’s localization is diffusive and it no-longer regulate adhesion morphology, which might contribute to the mis-regulated calcium signaling observed in cancer cells including stretch dependent apoptosis^16^.

There is growing evidence that covalent modifications in maturing adhesions alter adhesion proteins as well as possibly membrane proteins in normal cells^20,60^. The release of Ca^2+^ near adhesions contributes to protein processing by the adhesion signaling hub, which is needed for adhesion maturation and cell growth on a rigid fibronectin matrix possibly through a localized calpain-dependent pathway. The paxillin-associated calcium indicator shows that the Piezo1-dependent influxes of calcium are much more numerous than observed with the cytoplasmic calcium indicator. Thus, the localization of Piezo1 to the adhesions under force will provide a very localized activation of calpain and other calcium-dependent enzymes to modify adhesions and membrane proteins in specific locations.

In transformed cells, growth appears independent of local Ca^2+^ entry at rigid matrix sites and Piezo1 is decoupled from traction forces and adhesion signaling. It is commonly believed that transformed cells are related to wound healing where cell growth for repair is needed and localized signaling from matrices is not needed^61^. Further, in many cancers, the elevated growth rates correlate with high levels of calpain that could amplify small Ca^2+^ leaks to support the growth activity. This could help to explain how stretch could cause general changes in the plasma membrane activating Ca^2+^ entry through Piezo1 by local membrane tension or curvature changes that would cascade into an apoptotic process^16^. Most importantly, the general activation of growth in cancer cells appears to occur without localized signaling at adhesions by Piezo1.

In the larger picture of adhesion maturation, there are many adhesion proteins that bind to adhesions in a force-dependent manner^62,63^. It is surprising, however, that the inhibition of the release of many adhesion proteins by the inhibition of PTPN12 or expression of virulence bacterial protein IpaA did not block the release of Piezo1 upon myosin inhibition. Further, Piezo1 is strongly bound to the adhesions and has a lifetime similar to integrin beta3 in adhesions that depends on contractility. The stronger retention of Piezo1 at adhesions upon the transition from integrin beta3 to beta1 may aid in the stabilization of such adhesions for development of higher forces on matrix contacts through continued local influx of calcium^64^. The integrin subtype dependence of Piezo1 dynamics could be of great importance for the physiological function of Piezo1, e.g. in vascular development^4^. Piezo1 localization likely plays a critical role in processes such as vascular development and tumor angiogenesis where both Piezo1 and β3 integrins are indispensable. Piezo1 knock-out mice die as embryos due to vascular defects, and inactivation of beta3 inhibits the angiogenesis process. Mechanical forces such as shear forces are very important in these processes^4^. The force dependent co-localization and turnover of Piezo1 at β3 integrin adhesions could be synergistic with downstream events and provides a mechanism of tension-dependent vascular growth that is critical in development. Thus, we suggest that the contractility dependent recruitment of Piezo1 to focal adhesions is a novel crosstalk mechanism between the two major mechanotrasduction hubs of the cell that play critical roles in adhesion function and are lost in most tumor cells (Figure 8).

## Material and methods

### Cell culture and constructs

The mEmerald-integrin β3, mEmerald-integrin β1, paxillin-mapple, paxillin-BFP and talin-GFP were gifts from Michael Davidson (Addgene plasmid # 56330, #54129, #54935). The piezo1-GFP-1591 and piezo1-mruby3-1591 plasmids were generated as previously described^38^. The RhoAV14 construct was from Schwartz lab^65^. TPM2.1-YFP construct was from Gunning Lab^66^. The A431 ipa peptide plasmid was generated as previously described^67^. The paxillin calcium sensor was generated by PCR cloning of jGCaMP7s segment from pGP-CMV-jGCaMP7s plasmid from addgene^68^ and inserted into the N-terminal of a paxillin-mScarleti construct derived from paxillin-mapple.

siRNA for tropomyosin 2.1 was synthesized as previously described by Dharmacon^43^. siRNA for human Piezo1 was ordered from Dharmacon (ON-TARGETplus human Piezo1 siRNA, SMARTpool, L-020870-03-0010) and Merck Sigma-Aldrich (SASI_HS01_00208585) or Life Technologies(cat no #21816). RT-PCR analysis showed that both siRNA caused >90% knock down at the mRNA level two days after transfection. The siRNA were transfected using RNAiMAX reagent from ThermoFisher following manufacture’s protocols.

Human Foreskin fibroblast (HFF), HUMEC, HUVEC, HT1080, A2058, A375P, and MCF-10A cells were obtained from ATCC. MDA-MB-231, SKOV3, cells were from Ruby Huang’s lab. HEK293T cells with piezo1 KO were from Ardem Patapoutian’s lab. HFF, MDA-MB-231, HEK293T Piezo1 KO and cells were cultured in 1X DMEM supplemented with 5 mM sodium pyruvate and 10% of Hi-FBS. MCF-10A cells were cultured using fomulated media as reported previously^22^ and HUVEC cells were cultured in endothelial growth media from Sigma.

### Ca^2+^ influx measurements

For calcium level measurements using force sensors, cells were transfected with the GCAMP7S-paxillin-mScarleti construct and seeded on fibronectin coated glass bottom coverslips overnight. Transfected cells were imaged using TirfM microscrope using 488 and 561 laser lines. Specific band pass excitation/emission filter cubes (GFP-HQ and mCherry-HQ from Nikon) were used to avoid fluorescence crosstalk between channels. For comparison between different samples, all imaging parameters (laser power, tirf angle, camera exposures) were kept constant. Only cells with comparable calcium sensor expression levels (judging from fluorescence intensity from mScarlet channel) were picked for analysis to minimize potential bias due to sensor concentration differences.

The local Ca^2+^ influx of the cells was measured following Ellefsen et. al^26^. Briefly, the HFF cells were loaded with 4 uM Ca^2+^ indicator Cal-520-AM for 2 hours. Then the cells were detached using enzyme free dissociation media and resuspended in complete growth media. The cells were allowed to recover in suspension for 20 min before seeding on glass bottom coverslips coated with fibronectin. After 2 hours the buffer was switched to Ringer’s balanced salt and proceeded for imaging on Nikon Tirf system with pulsed laser lines to minimize photodamage of the sample^69^. The calcium images of individuals cells were acquired at 100Hz for 5000 frames. The deltaF/F images of the cells were generated using a custom matlab script to reveal individual Ca^2+^ influx events. Such events were tracked using the Trackmate ImageJ plugin^70^.

### Cell spreading assay

In the cell spreading assay, cells were transfected with respective plasmids 36-48 hours before experiments using Neon transfection system (Thermo Fisher) or JetOptimus chemical transfection reagent (PolyPlus) following manufacturer’s recommendations. On the day of experiment, the cells were detached using an enzyme-free cell detaching buffer (Cell Dissociation Buffer, enzyme-free, PBS, Gibco), spun-down and resuspended in complete media (1X DMEM, 10% FBS). The suspended cells were allowed to recover by incubating in the cell culture hood for 20 min and then seeded onto glass bottom Petri dishes coated with ECM molecules of interest (10 ug/ml collagen I, 10 ug/ml fibronectin, 10 ug/ml VTN-N, and 10 ug/ml laminin for 1 hour). Imaging was done on a Nikon Tirf system with 405,488,561 and 640 laser lines. The cell spreading area and adhesion size were analyzed automatically using CellProfiler^71^.

### Immunostaining and antibodies

Cells of interest seeded on glass bottom Petri dishes were fixed with 4% paraformaldehyde (PFA) for 30 min at 37 degrees and washed with 1X PBS three times. The fixed cells were permeabilized/blocked by incubation with TBST (1X TBS, 0.1% Tween-20) containing 5% Bovine Serum Abumin (BSA) overnight at 4 degrees. Primary antibodies were diluted in TBST, 5% BSA solution according to manufactures’ recommendations and added to the cells overnight at 4 degrees. The stained cells were washed with 1X PBS and stained with secondary antibody for 1 hour at room temperature.

Polyclonal Rabbit and monoclonal mouse anti-Piezo1 was from Novus Biologicals (NBP1-78446, NBP2-75617), both have been validated by Piezo1 knock down/knock off experiments^15,72^. For all Piezo1 localization quantifications, the monoclonal NBP2-75617 antibody was used. Mouse anti-Paxillin antibody was from Merck Millipore (MAB 3060), Rabbit anti-paxillin antibody was from abcam(ab32115). Mouse anti-activated integrin β1 antibody was from BD Biosciences (9EG7). Rat anti integrin αvβ3 was from Millipore(LM609).

### FRAP analysis

FRAP analyses were carried out on an ILAS2 TIRF image system similar to reported previously. Briefly, HFF, HT1080, or MDA-MB-231 cells transiently co-transfected with Piezo1-GFP and Paxillin-mapple, or Piezo1-mruby3 and mEmerald-integrin β3 were seeded on Fibronectin coated glass bottom dishes overnight. 2-5 square-shaped regions of interests (∼3 um length) covering focal adhesions were selected for each cell and bleached. The recovery of fluorescence in both fluorescent channels was observed for from 360 to 600 seconds after photobleaching. The recovery curves were normalized by dividing by average intensity of each ROI before bleaching and the average and standard deviation of the recovery were calculated.

## Supplementary Figures

**Figure S1.**
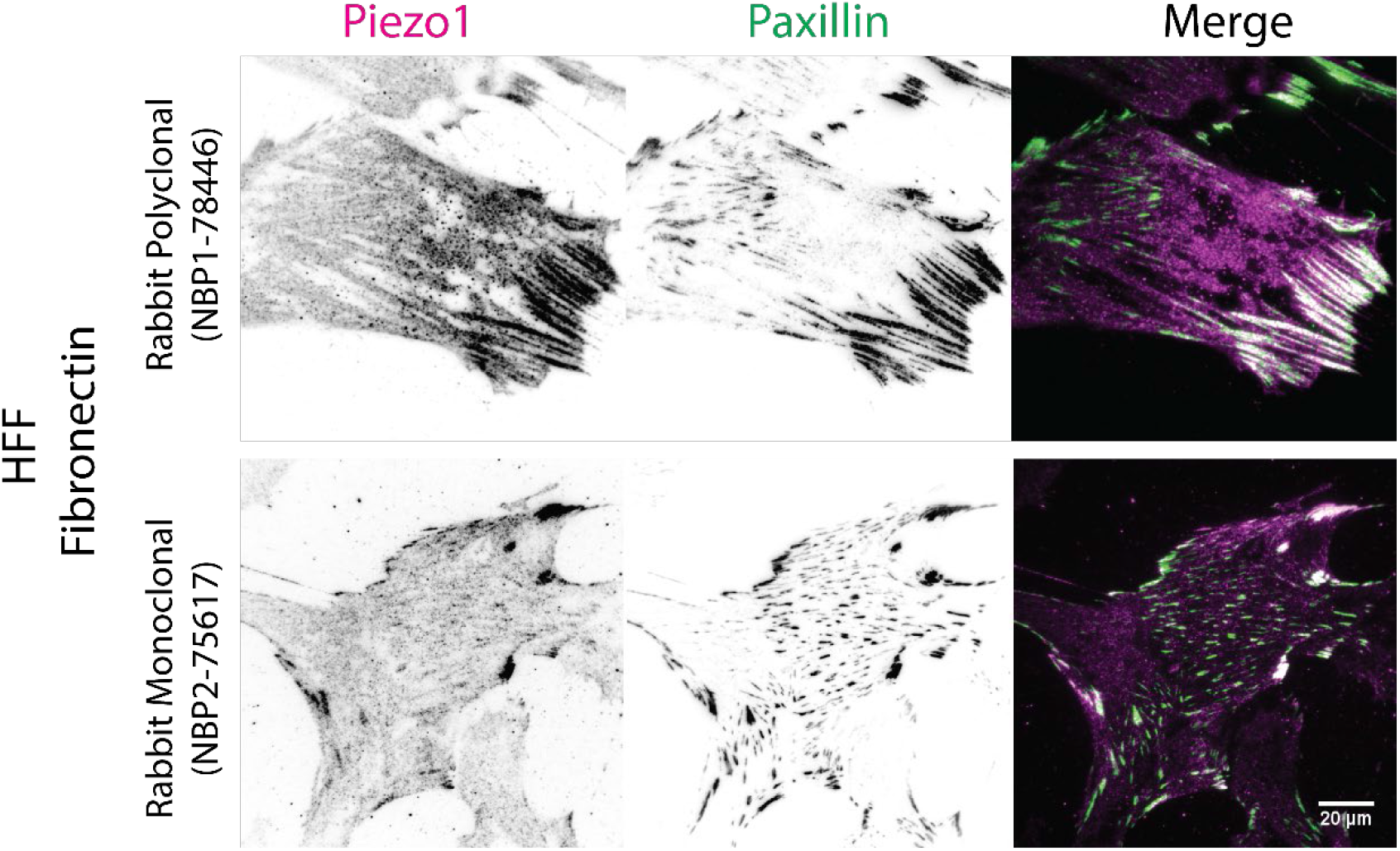
Immunostaining of Piezo1 localization of HFF cells. HFF cells were seeded on fibronectin surfaces overnight, fixed with paraformaldehyde and stained for Piezo1 with two validated Piezo1 antibody and paxillin.

**Figure S2.**
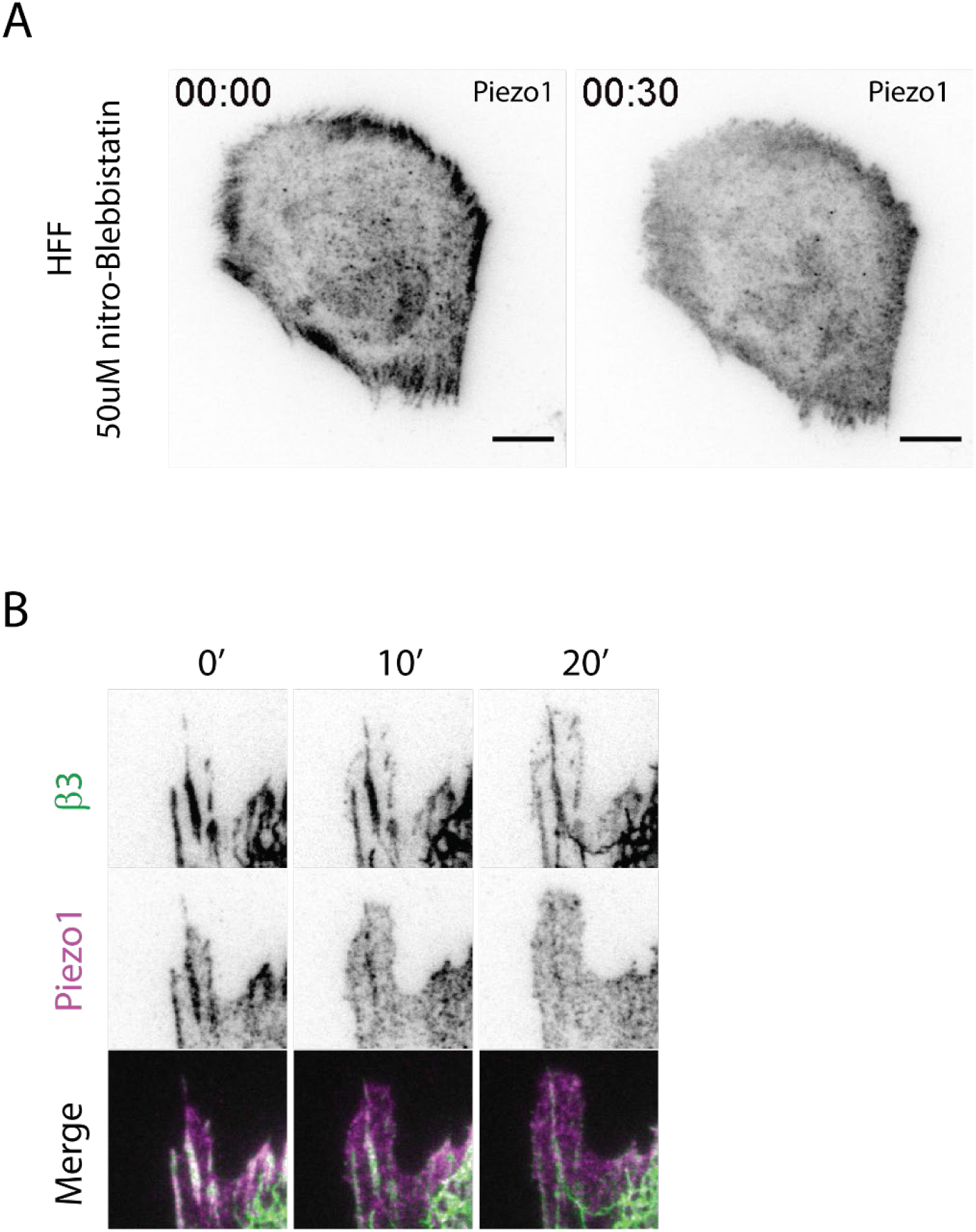
(A) HFF cells transiently transfected with Piezo1-mruby3 were treated with 50 uM nitro-Blebbistatin at time 0. After 30 min incubation, the localization of piezo1 at focal adhesions were lost. Scale Bars denote 10 um. (B) HFF cells transiently transfected with mEmerald-integrin β3 and Piezo1-mruby3 were pre-treated with 2.5 uM PTP-PEST inhibitor that inhibits adhesion turnover by phosphatases before addition of 20 uM Y-27632. In this condition, Piezo1 dissociated from adhesions before integrin β3 upon tension release.

**Figure S3.**
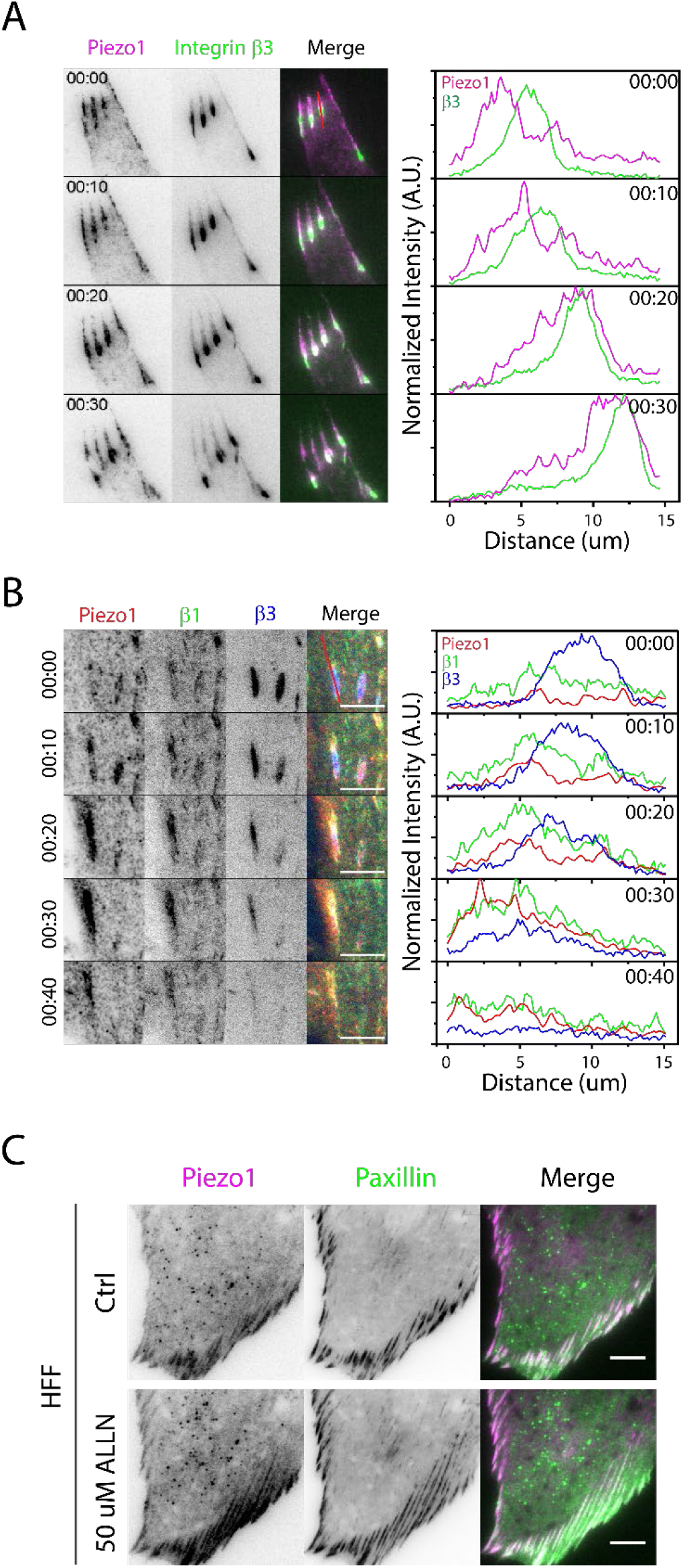
Dynamics of Piezo1 localization in relation to integrins. (A) Adhesion time lapse HFF cells transiently transfected with mEmerald-integrin β3 and Piezo1-mruby3. The right panel showed normalized intensity profiles of the adhesions marked by red line over time. (B) Adhesion time lapse of HFF cells transfected with BFP-integrin β3, mEmerald-integrin β1 and Piezo1-mruby3. The right panel showed normalized intensity profiles of the adhesions marked by red line over time. The normalizations were done by dividing the intensity profiles by the maximum intensity value observed for each protein in the whole time series. Scale Bars denote 10 μm.. (C) Elongation of Piezo1 containing adhesions upon calpain inhibition. Live imaging of HFF cells before and four hours after treatment with 50 μM ALLN. The adhesions were marked by paxillin-mapple. Scale Bars denote 10 um.

**Figure S4.**
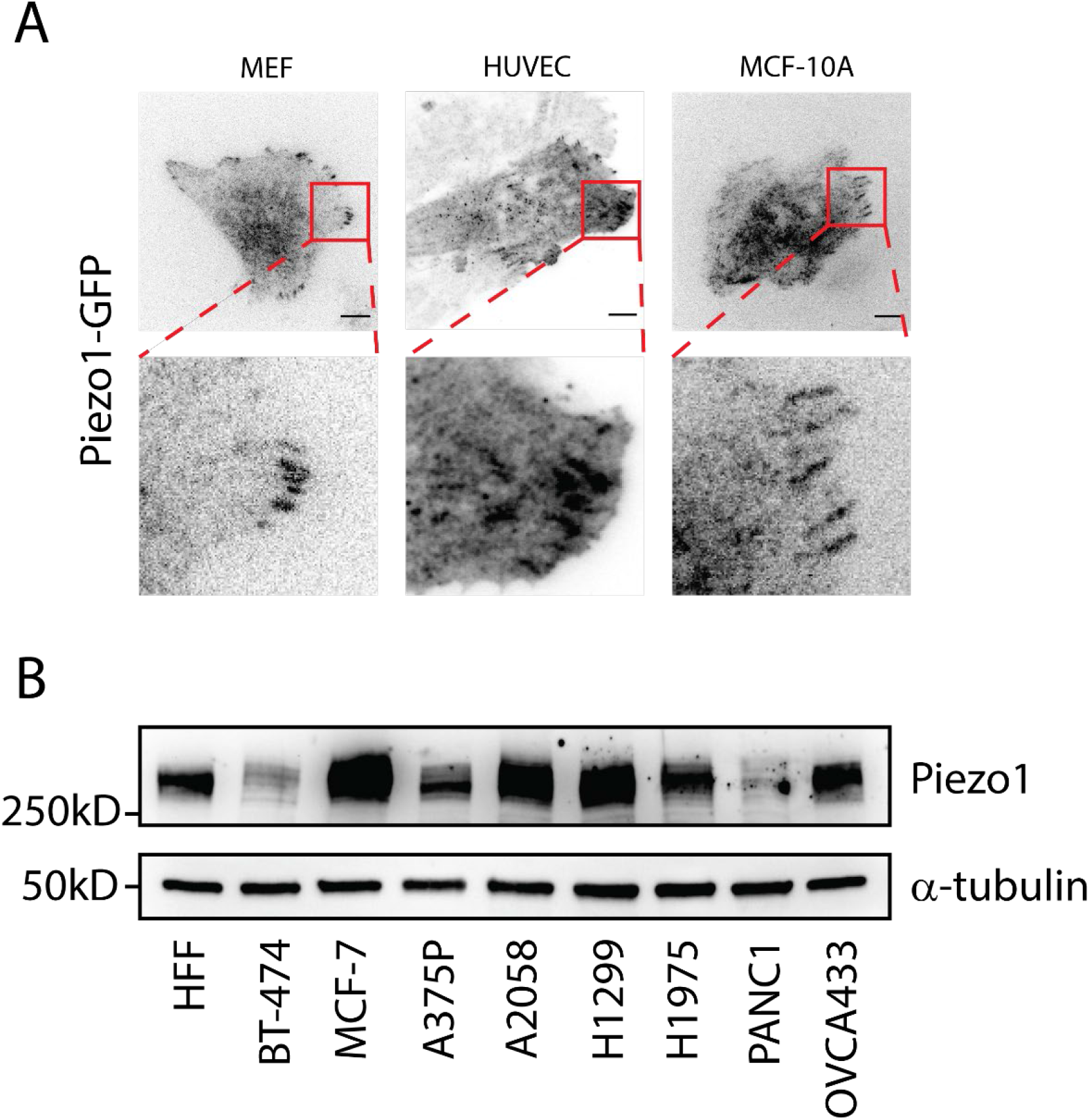
(A) TIRF live images of Piezo1-GFP localization in more normal cell lines tested. (B) Western blot images of Piezo1 expression in various cancer cell lines compared against HFF cells.

**Figure S5.**
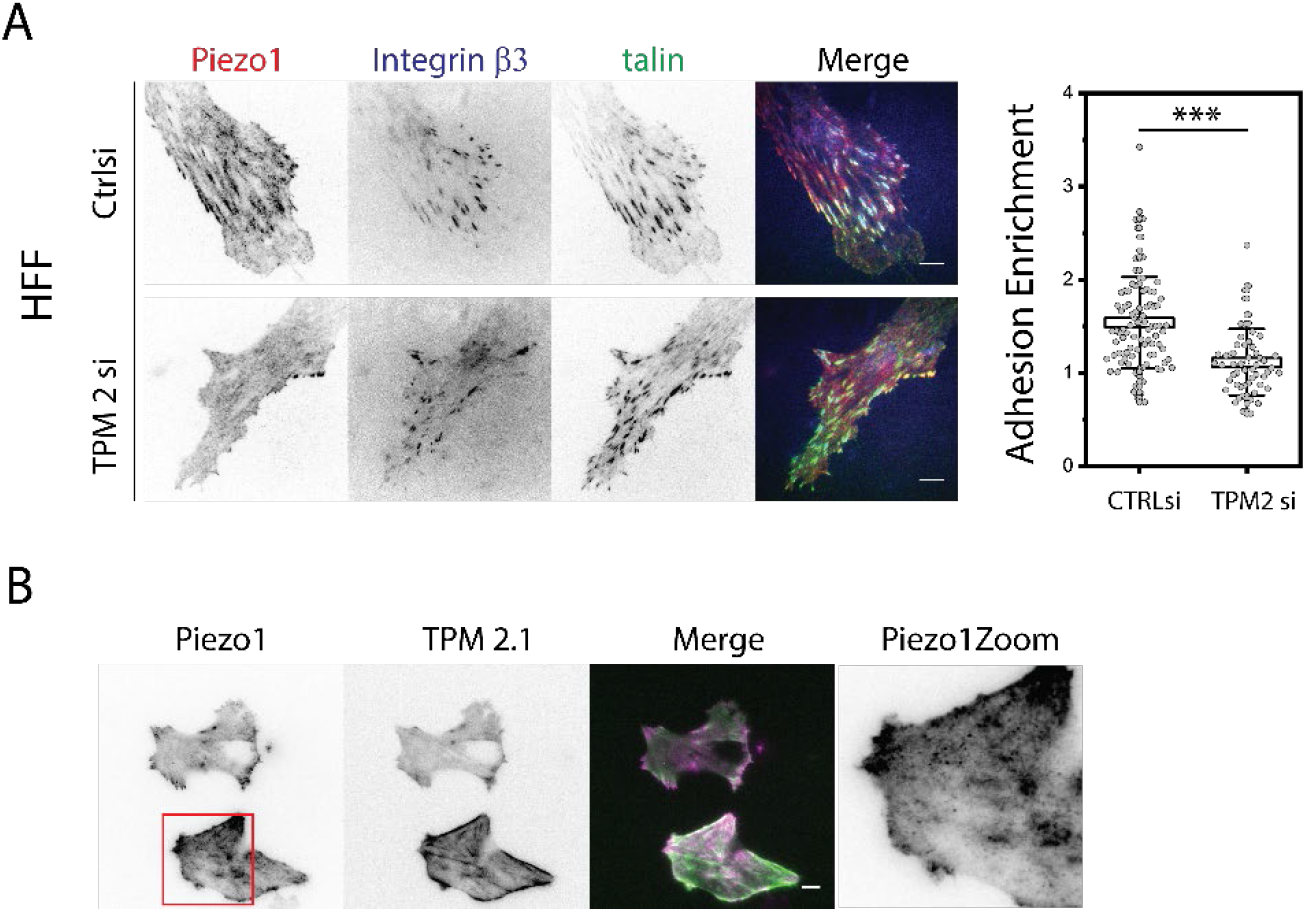
Piezo1 localization in cells with altered transformation state. (A) Representative HFF cells with Control siRNA and TPM 2 siRNA knock down were transfected with Piezo1-mruby3, Integrin β3-BFP and talin-GFP. The cells were seeded on fibronectin surfaces overnight and imaged by TIRFM. Right Panel: quantification of adhesion enrichment with Ctrlsi an d TPM2 si (C) Transient expression of TPM 2.1 -YFP in MDA-MB-231 cells led to the recruitment of Piezo1 to the end of stress fibres decorated by TPM 2.1 (TOP panel). The images present representative cells out of > 20 imaged. Scale Bars denote 10 um.

**Figure S6.**
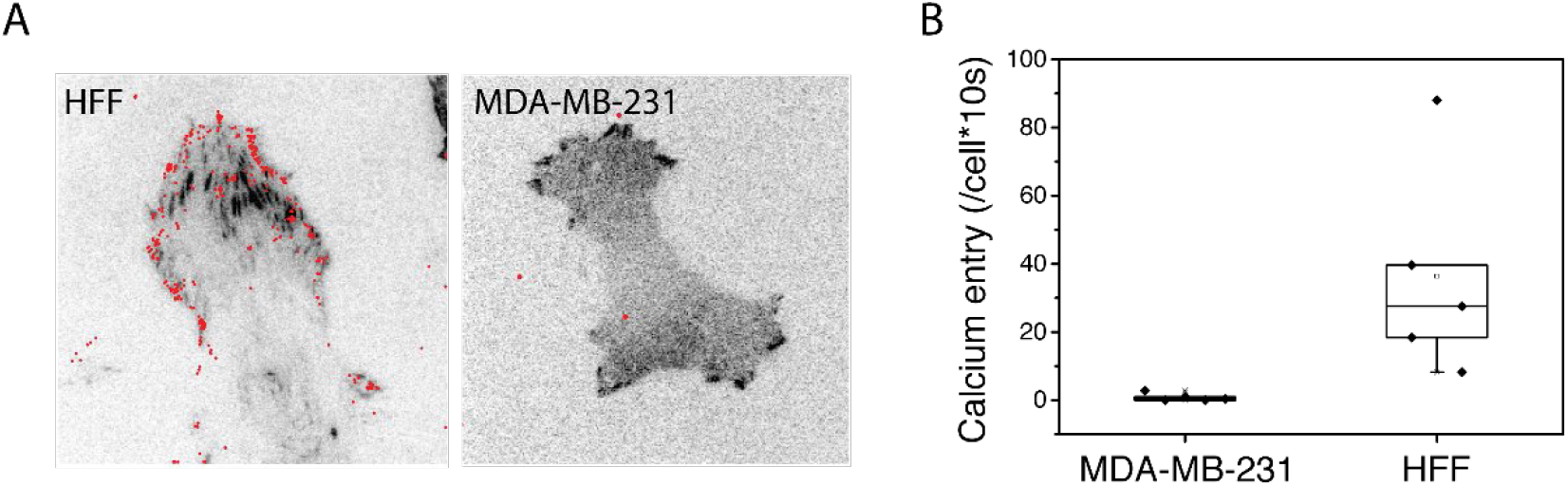
HFF and MDA-MB-231 cells showed distinct local calcium entry pattern. (A) Locations of local calcium entry events (marked by red dots) in HFF cells and MDA-MB-231 cells measured at 100 frames/s for 50s, overlaid with focal adhesions marked by paxillin. Significantly more local calcium signal is observed in HFF cells. (B). Box plot of the rate of local calcium events per cell for HFF and MDA-MB-231 cells. Each point denotes an independent cell.

**Figure S7.**
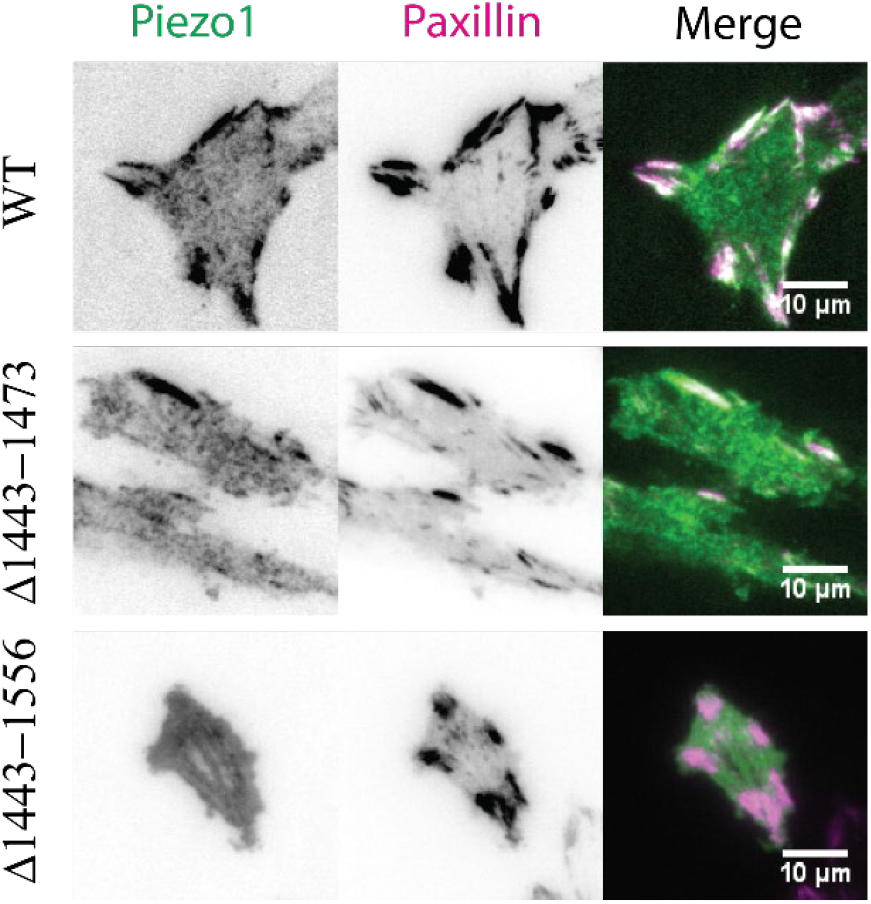
Linker domain mutants of Piezo1. Representative images of FAK-/- MEF cells transfected with paxillin-mapple and wild type/linker deletion mutant of Piezo1. Similar behavior were observed in 3 biological repeats with > 10 cells for each condition.

